# The requirement for the amino acid co-germinant during *C. difficile* spore germination is influenced by mutations in *yabG* and *cspA*

**DOI:** 10.1101/427021

**Authors:** Ritu Shrestha, Joseph A. Sorg

## Abstract

*Clostridium difficile* spore germination is critical for the transmission of disease. *C. difficile* spores germinate in response to cholic acid derivatives, such as taurocholate (TA), and amino acids, such as glycine or alanine. Although the bile acid germinant receptor is known, the amino acid germinant receptor has remained elusive. Here, we used EMS mutagenesis to generate mutants with altered requirements for the amino acid co-germinant, similar to the strategy used previously to identify the bile acid receptor, CspC. Surprisingly, we identified strains that do not require amino acids as co-germinants, and the mutant spores germinated in response to TA alone. Upon sequencing these mutants, we identified different mutations in *yabG.* In *C. difficile, yabG* expression is required for the processing of CspBA to CspB and CspA and preproSleC to proSleC during spore formation. A defined *yabG* mutant exacerbated the EMS mutant phenotype. Moreover, we found that various mutations in *cspA* caused spores to germinate in the presence of TA alone without the requirement of an amino acid. Thus, our study provides evidence that apart from regulating the CspC levels in the spore, CspA is important for recognition of amino acids as co-germinants during *C. difficile* spore germination and that two pseudoproteases (CspC and CspA) function as the *C. difficile* germinant receptors.

## Introduction

*Clostridioides difficile* (formerly *Clostridium difficile*) [1–3] is a Gram-positive, spore-forming pathogenic bacterium, and has become a leading cause of nosocomial diarrhea in the United States [4, 5]. *C. difficile* infection (CDI) is commonly the result of disruption to the gut microflora caused by antibiotic use [5–7]. Due to the broad-spectrum nature of many antibiotics, alterations to the ecology of the colonic microbiome results in the loss of the colonization resistance that is provided by the microbiota. Subsequently, patients are treated with other, broad-spectrum, antibiotics (*e.g.,* vancomycin or fidaxomicin) which treat the actively growing, toxin-producing, vegetative cells [8]. Although these antibiotics alleviate the primary symptoms of disease, the continued disruption to the colonic microbiome results in frequent CDI recurrence. The symptoms of CDI are caused by the actions of two secreted toxins. TcdA (an enterotoxin) and TcdB (a cytotoxin) are endocytosed by the colonic epithelium and inactivate the Rho-family of small GTPases leading to loss of barrier function and inflammation of the colonic epithelium [7].

Though *C. difficile* vegetative cells produce the toxins that cause CDI, they are strictly anaerobic and only survive short periods of time outside the anaerobic colonic environment [9]. However, the spores that are produced by the vegetative form are critical for transmission between hosts because of their resistance to environmental factors such as heat, UV, chemicals and, importantly, oxygen [10–14]. The overall architecture of *C. difficile* spores is conserved among all endospore-forming bacteria. The centrally-located core is composed of DNA, RNA, ribosomes and proteins necessary for the outgrowth of a vegetative cell, post germination [11, 14]. The DNA in the core is protected from UV damage by small acid soluble proteins (SASPs) and much of the water in the core is replaced by pyridine-2, 6-dicarboxylic acid (dipicolinic acid; DPA), chelated with calcium (CaDPA), which provides heat resistance to the spores [11, 14, 15]. The core is surrounded by an inner membrane composed of phospholipids with minimal permeability to small molecules, including water [15]. A thin germ-cell-wall layer surrounds the inner membrane and becomes the cell wall of the vegetative cell upon outgrowth. A thick layer of specialized peptidoglycan (cortex) surrounds the germ cell wall and helps constrain the core against osmolysis [15]. Finally, surrounding the cortex is the outer membrane which, initially, serves as an organization structure / point for the coat layers but may be lost later during spore development [11, 16–20]. All these features of endospores contribute to ensuring that the spores remain metabolically dormant.

Though dormant, spores still sense their environment for species-specific germination-inducing small molecules and, when appropriate germinants are present, initiate the process of spore germination. Much of our knowledge of spore germination comes from studies in *Bacillus subtilis* (a model organism for studying spore formation and germination). In *B. subtilis,* and most other endospore-forming bacteria, germination is activated upon binding of the germinants to the Ger-type germinant receptors that are deposited in or on the inner spore membrane [21, 22]. In *B. subtilis,* this event triggers an irreversible process whereby CaDPA is released through a channel composed of the SpoVA proteins [23–29]. The release of CaDPA is an essential step for the germination process because it results in the rehydration of the spore core and permits the eventual resumption of metabolic activity. In *B. subtilis,* the cortex peptidoglycan layer is then degraded, and this event can be activated by the CaDPA that is released from the core [30].

In contrast to *B. subtilis, C. difficile* does not encode orthologues of the Ger-type germinant receptors suggesting that *C. difficile* spore germination is fundamentally different than what occurs in most other studied organisms [10]. Germination by *C. difficile* spores is triggered in response to certain bile acids in combination with certain amino acids [10, 31, 32]. In all identified *C. difficile* isolates, the cholic acid-derivative, taurocholate (TA), is the most efficient bile acid at promoting spore germination, and glycine is the most efficient amino acid co-germinant [33–35]. Recently, calcium was identified as an important contributor to spore germination, though its role remains unclear [36]. Calcium may function as a bona fide spore co-germinant, a co-factor for a process essential for spore germination and / or as an enhancer for amino acids [13, 36].

In a screen to identify ethyl methane sulfonate (EMS) generated mutants that could not respond to TA as a spore germinant, we identified the germination-specific, subtilisin-like, pseudoprotease, CspC, as the *C. difficile* bile acid germinant receptor [37]. Prior to work performed in *C. difficile,* the Csp locus was best studied in *Clostridium perfringens* [38–41]. *C. perfringens* encodes three Csp proteases, CspB, CspA, and CspC, that are predicted to cleave the inhibitory pro-peptide from proSleC, a spore cortex hydrolase that degrades the cortex peptidoglycan, thereby activating the protein [38–41]. In *C. difficile,* CspB and CspA are encoded by one ORF and the resulting protein is post-translationally processed into CspB and CspA and then further processed by the sporulation specific protease, YabG [14, 42–44]. CspC is encoded downstream of *cspBA* and is part of the same transcriptional unit. Interestingly, the catalytic residues in CspA and CspC are lost, suggesting that only the CspB protein can process proSleC to its active, cortex degrading form [37, 44]. Although present in the spore, and essential for *C. difficile* spore germination, the CspA pseudoprotease has only been shown to regulate the incorporation of CspC into the spore [42, 43].

In our working model for spore germination, activation of CspC by TA leads to the activation of the CspB protein which cleaves proSleC into its active form. Activated SleC degrades cortex and the core releases CaDPA in exchange for water by a mechanosensing mechanism [45]. Because the receptor with which amino acids interact is unknown, we sought to screen for chemically-generated mutants that have altered amino acid requirements during spore germination [similar to the strategy used to identify the bile acid germinant receptor [37]]. Here, we report that a mutation in *yabG* gene results in strains whose spores no longer respond to or require amino acid co-germinants but respond to TA alone as a spore germinant. We hypothesize that the misprocessing of CspBA in the *yabG* mutant leads to this phenotype and provide evidence that short deletions in CspA alter the requirements for a co-germinant during spore germination. Our results suggest that CspA is the amino acid co-germinant receptor.

## Materials and Methods

### Growth conditions

*C. difficile* strains were grown on BHIS agar medium (Brain heart infusion supplemented with 5 g / L yeast extract and 1 g / L L-cysteine) in an anaerobic chamber (Model B, Coy Laboratories Grass Lake, MI) at 37 °C (85% N_2_, 10% H_2_, and 5% CO_2_). Antibiotics were added as needed (15 μg / mL of thiamphenicol, 10 μg / mL lincomycin, and 20 μg / mL uracil). Deletion mutants were selected on *C. difficile* minimal medium (CDMM) supplemented with 5 μg / mL 5-fluoroorotic acid (FOA) and 20 μg / mL uracil. *Bacillus subtilis* was used as a conjugal donor strain to transfer plasmids into *C. difficile* and was grown on LB medium with 5 μg / mL of tetracycline and 2.5 μg / mL of chloramphenicol. Conjugation was performed on TY medium (3% Bacto Trypone, 2% yeast extract, and 0.1% thioglycolate) with or without uracil. *E. coli* DH5α and *E. coli* MB3436 were grown on LB medium supplemented with 20 μg / mL chloramphenicol at 37 °C.

### Spore purification

*C. difficile* spores were purified as described previously [32, 33, 45, 46]. Briefly, the strains without plasmids were grown on BHIS agar medium while strains with plasmid were grown on BHIS agar medium supplemented with 5 μg / mL thiamphenicol and allowed to grow for 4 days. Cells from each plate were scraped into 1 mL sterile water and incubated at 4 °C overnight. Next, the cells were washed five times with water and combined into 2 mL total volume. The washed spores were layered on top of 8 mL of 60% sucrose and centrifuged at 4,000 X g for 20 minutes. The supernatant was discarded, and the spores were washed five more times with water and incubated at 4 °C until use.

### EMS mutagenesis and phenotype enrichment

An overnight culture of wild type *C. difficile* UK1 vegetative cells were diluted to an OD_600_ of 0.05 into two, separate 15 mL falcon tubes containing BHIS liquid and grown for 3 - 4 hrs. To one of the cultures, ethyl methane sulfonate (EMS) was added to a final concentration of 50 μg / mL; the other culture was untreated for use as a negative control. The cultures were grown for 3 hours and then centrifuged at 3,000 X g for 10 min. The supernatants were discarded, and pellets were washed two more times with BHIS. After the final wash, the pellets were suspended in 40 mL BHIS medium and allowed to recover overnight. The recovered EMS-treated cells were then plated onto 10 - 12 BHIS plates (25 μL on each plate) to produce spores. Spores were purified from the EMS-treated strain as described above. Purified spores were heat activated at 65 °C for 30 minutes before enrichment as shown in Figure 1A. The EMS-treated spores were treated for 15 minutes with HEPES buffer (50 mM HEPES, 100 mM NaCl, pH 7.5) supplemented with 10 mM TA and 10 mM betaine and then washed twice with buffer. Germinated, washed spores were plated onto BHIS agar medium for spore formation. This procedure was repeated iteratively for 4 – 5 times before isolating candidate strains for phenotypic screening.

**Figure 1.**
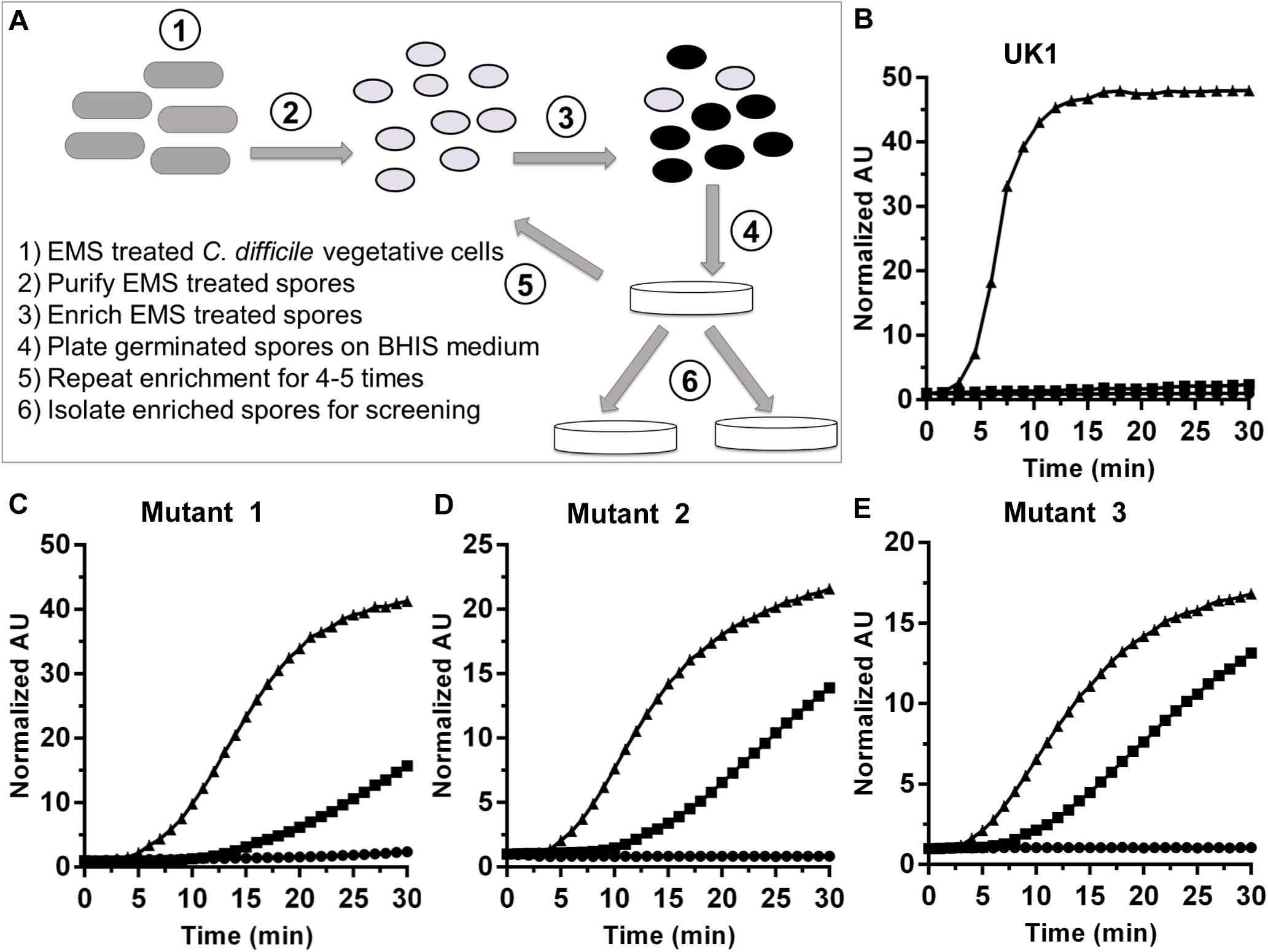
EMS mutagenesis generates *C. difficile* strains with altered requirements for the amino acid co-germinant. (A) Schematic of the EMS mutagenesis strategy where (1) *C. difficile* UK1 vegetative cells were treated with EMS, (2) EMS mutagenized cells were recovered and allowed to form spores, (3) purified spores were suspended for 15 minutes in germination buffer supplemented with 10 mM TA and 10 mM betaine and then washed twice with buffer alone, (4) germinated spores were plated onto BHIS agar medium to permit colony formation, (5) strains were enriched by exposure TA and betaine, iteratively, 4 – 5 times, and (6) spores from the mutant strains were isolated for screening. The germination phenotype of wild type *C. difficile* UK1 (B) and mutant spores (C – E) were screened by measuring CaDPA release in presence of (●) 30 mM glycine, (■) 10 mM TA or (▲) 10 mM TA and 30 mM glycine. The data points represent the data from a single germination experiment. Germination plots performed in triplicate yielded error bars that obscure the data. For transparency, all germination plots can be found in Figure S1.

### Characterizing mutant phenotypes

The mutant spores were initially characterized by measuring the CaDPA release from germinating spores. Spores were heat activated at 65 °C for 30 minutes and suspended in water at an OD_600_ = 50. The spores were then added to final OD of 0.5 in 100 μL final volume of HEPES buffer containing 250 μM Tb^3^+, 10 mM TA and / or 10 mM betaine in a 96 well plate. The CaDPA release was measured using a SpectraMax M3 plate reader for 30 minutes at 37 °C with excitation at 270 nM and emission at 545 nM with a 420 nM cutoff.

The phenotype of the *yabG* mutant (RS08) and *cspBA* deletion mutants were determined by measuring changes to the OD_600_ and release of DPA. OD_600_ was monitored at 37 °C for 1 – 2 hrs using a plate reader. The spores were added to a final OD of 0.5 in HEPES buffer supplemented with 30 mM glycine alone or 10 mM TA alone or 10 mM TA and 30 mM glycine in 100 μL final volume. CaDPA release was measured as described above with spores containing final OD_600_ of 0.25.

### DNA Sequencing

High-quality genomic DNA was purified from logarithmically growing *C. difficile* cells [47]. Genomic DNA from the EMS-mutant strains and the wild type parent was sent for paired end sequencing at Tufts University Genomics Core. Reads were aligned to the R20291 genome using DNASTAR software and SNPs determined (DNASTAR, Madison, WI). RAW sequence (fastq) reads were uploaded to the NCBI Sequence Read Archive as follows (Table 1): *C. difficile* UK1 parent (SRS3677310), Mutant 20C (SRS3677309), Mutant 27E (SRS3677307), Mutant 30A (SRS3677308), Mutant 30C (SRS3677311), Mutant 31D (SRS3677312).

**Table 1.** Location of the mutations in *yabG* found in the EMS mutant strains.

| Mutant | Reference | Coding region | Promoter region |
| --- | --- | --- | --- |
| 20C<br>(Mutant 1) | 4048886 | G-T(Ala46Asp) |  |
| 27E<br>(Mutant 2) | 4049063 |  | G-42A |
| 30A<br>(Mutant 3) | 4048565 | G-A<br>(Pro153Leu) |  |
| 30C | 4048913 | C –T<br>(Gly37Glu) |  |
| 31D | 4049030 |  | C-8T |

Nucleotide positions in the promoter region are mapped from the ATG of the open reading frame (the *yabG* transcriptional start site is unknown)

### Molecular Biology

The oligonucleotides used for making the strains and plasmids used in this study are listed in Table S1. *yabG* and *sleC* mutants were created in the *C. difficile* R20291 background by using the TargeTron mutagenesis system. The potential insertion sites for targeting the group II intron were found using the Targetronics algorithm (Targetronics, LLC.) and a gBlock (Integrated DNA Technologies, San Jose, CA) of the group II intron targeted to *yabG* at the 279^th^ nucleotide, relative to the start codon, was ordered (Table S1) and cloned into pJS107 at the HinDIII and BsrGI restriction sites using Gibson assembly. The ligation was then transformed into *E. coli* DH5α. The site to engineer a TargeTron insertion in *sleC* (pCA6) was previously described for *C. difficile* UK1 [46] and the same protocol was used to engineer the insertion into R20291. The TargeTron insertion plasmids were isolated from *E. coli* DH5α and transformed into *E. coli* MB3436 (a *recA*^+^ strain) and then into *B. subtilis* (BS49). Conjugation between *B. subtilis* and *C. difficile* R20291 was performed on TY-agar medium for 24 hrs before plating onto selection plates. Once the plasmid was inserted into *C. difficile,* the colonies were screened for tetracycline-sensitive and thiamphenicol-resistant colonies and confirmed with PCR. *yabG* or *sleC* TargeTron mutants were selected by plating the colonies onto BHIS supplemented with lincomycin. Lincomycin-resistant colonies were tested by PCR and confirmed by sequencing the mutation. *yabG* and *sleC* TargeTron mutants were renamed as RS08 and RS10, respectively (Table 3).

**Table 2.** Plasmid list with primer pairs to make the plasmids

| Plasmid | Description | Oligonucleotides<br>used (Supplemental | Reference |
| --- | --- | --- | --- |
|  |  | Table 1) |  |
| pRS92 | *yabG* G-block cloned into Zero blunt | 813 | This study |
| pRS93 | Plasmid to make TT insertion in *yabG* |  | This study |
| pRS97 | *yabG* Complementing plasmid | 906/907, 924/925 | This study |
| pCA6 | Plasmid to make TT insertion in *sleC* |  | [46] |
| pMTLYN4 | Plasmid to make deletion in *C. difficile* using allelic exchange |  | [61] |
| pJS107 | TargetTron vector |  | [37] |
| pJS116 | Empty vector |  | [37] |
| pJS165 | Tn916Ori cloned into pMTLYN4 to use *B. subtilis* conjugation for gene insertion | 207, 208 | This study |
| pRS110 | pJS116 with $\Delta 542-567$ in CspBA | 578/1232, 1233/1056 | This study |
| pRS111 | pJS116 with $\Delta 537-571$ in CspBA | 578/1234, 1235/1056 | This study |
| pRS112 | pJS116 with $\Delta 560-586$ in CspBA | 578/1236, 1237/1056 | This study |
| pRS113 | pJS116 with $\Delta 522-548$ in CspBA | 578/1238, 1239/1056 | This study |
| pRS114 | pJS165 with 1Kb upstream and 1Kb downstream of $\Delta 542-567$ in CspBA | 1160, 1163 | This study |
| pRS115 | Plasmid to make $\Delta 537-571$ in CspBA cloned into pJS165 | 1160, 1163 | This study |
| pRS116 | Plasmid to make $\Delta 560-586$ in CspBA cloned into pJS165 | 1160, 1163 | This study |
| pRS117 | Plasmid to make $\Delta 522-548$ in CspBA cloned into pJS165 | 1160, 1163 | This study |
| pMTLYN2C | Plasmid to restore *pyrE* |  | [61] |
| pRS107 | Tn916Ori cloned into pMTLYN2C to use *Bacillus subtilis* for restoring *pyrE* | 207, 208 | This study |
| pRS118 | Plasmid to make $\Delta 560-610$ deletion in CspA | 1160/1287, 1163/1288 | This study |
| pRS120 | Plasmid to make SRQS deletion in CspA | 1160/1329, 1163/1330 | This study |
| pRS121 | Plasmid to make SRQS deletion in preproSleC | 1361/1362, 1363/1364 | This study |

**Table 3.** Strain list

| *C. difficile* Strain | Description | Plasmids used | Reference |
| --- | --- | --- | --- |
| UK1 | Wild type, ribotype 027 |  | [62] |
| R20291 | Wild type, ribotype 027 |  | [63] |
| CRG2359 | R20291 $\Delta pyrE$ | | [61] |
| RS08 | *yabG::ermB* in R20291 background | pRS93 | This study |
| RS10 | *sleC::ermB* in R20291 background | pCA6 | This study |
| RS19 | CRG2359 with restored *pyrE* | pRS107 | This study |
| RS20 | CRG2359 with Δ522-548 deletion in CspB | pRS117 | This study |
| RS21 | CRG2359 with Δ560-586 deletion in CspA | pRS116 | This study |
| RS23 | CRG2359 with Δ542-567 in CspBA | pRS114 | This study |
| RS24 | CRG2359 with Δ537-571 in CspBA | pRS115 | This study |
| RS26 | RS21 with restored *pyrE* | pRS107 | This study |
| RS27 | CRG2359 with Δ560-610 in CspA | pRS118 | This study |
| RS29 | ΔSRQS (Δ580-584) in CspA | pRS120 | This study |
| RS30 | ΔSRQS (Δ117-121) in preproSleC | pRS121 | This study |
| RS31 | ΔSRQS in CspA and preproSleC | pRS120, pRS121 | This study |
| **Other Strains** |  |  |  |
| *E. coli* DH5α | Cloning vector |  | [64] |
| *E. coli* BL21(DE3) | Expression vector |  | [65] |
| *E. coli* MB3436 | *recA*<sup>+</sup> *E. coli* strain |  | Gift from Dr. Michael Benedik |
| *B. subtilis* Bs49 | Tn916 donor strain, Tet<sup>R</sup> |  | [66] |

A *yabG* complementing plasmid (pRS97) was created using primers 5’ XbaI_Prom_YabG and 3’ YabG_XhoI to amplify the *yabG* promoter and *yabG* coding regions and cloned into pJS116. The complementing plasmid was then inserted into *C. difficile* RS08 by conjugation with *B. subtilis*, as described above.

To engineer the required site-specific deletions, the *pyrE*-mediated allelic exchange strategy was used with the *C. difficile* CRG2359 strain (R20291 Δ*pyrE*). Briefly, 1 kb upstream and 1 kb downstream fragments that surround the desired mutation in *cspBA* or *preprosleC* were cloned into pJS165 using primers listed in Table S1. The plasmids were inserted into *C. difficile* CRG2359 strain using *B. subtilis* conjugation as described above. The strains containing the plasmids were then passaged several times to encourage the formation of single recombinants before passing onto CDMM-FOA-uracil medium. Thiamphenicol-sensitive candidate strains were tested by PCR for the desired mutations and confirmed by sequencing for the mutagenized regions. Where indicated, *pyrE* was restored to wild type using pRS107 (Table 2).

### Western blot

Samples were prepared for CspB, CspA, and CspC western blot by extracting soluble proteins from 2 × 10^9^ / mL spores [R20291, *yabG::ermB, yabG::ermB* pRS97 (pyabG)]. For the protein standard, recombinant CspB, SleC and CspC protein were purified using a previously described protocol [33]. Standard amount of protein or number of spores were solubilized in NuPAGE sample buffer (Life Technologies) and heated at 95 °C for 20 minutes. Equal volume of spore extracts and recombinant CspB, CspC or SleC standard proteins were separated by SDS-PAGE. Proteins were then transferred onto low-fluorescence polyvinylidene difluoride membrane (PVDF) at 30V for 16 hours. The membranes were then blocked in 10% skimmed milk in TBS (Tris-buffered saline) and washed thrice with TBS containing 0.1% (vol / vol) Tween-20 (TSBT) for 20 minutes each at room temperature. The membranes were then incubated with anti-CspB, anti-CspC or anti-SleC antibodies for 2 hours and washed with TSBT thrice. For the secondary antibody, AlexaFlour 555-labeled donkey anti-rabbit antibody was used to label the membranes for 2 hours, in the dark. The membranes were washed again, thrice, with TBST, in the dark, and scanned with GE Typhoon Scanner using Cy3 setting, an appropriate wavelength for the Alexa Flour 555 fluorophore. The fluorescent bands were quantified using ImageQant TL 7.0 image analysis software. Intensity of the extracted protein in each blot was compared to the standard curve that was generated from the recombinant protein included on each blot.

To analyze SleC activation, equal number of spores were suspended in HEPES buffer supplemented with 30 mM glycine or 10 mM TA or 10 TA and 30 mM glycine and incubated at 37 °C for 1 hr to 2 hrs. The samples were then centrifuged at 15,000 X g for 1 minute and pellets were suspended in NuPAGE sample buffer and heated for 20 min at 95 °C. The suspension was centrifuged at max rpm for 10 min. The supernatant was separated and transferred into new tubes. The samples were stored at −20 °C until use. For CspC, CspA and CspB western blots, an equal number of spores were suspended in HEPES buffer and boiled to extract the protein and loaded in 10% SDS PAGE gel. The spore extracts were then transferred into nitrocellulose membrane for western blot analysis.

### Statistical Analysis

All germination assays were performed in triplicate and data points represent the average of three independent experiments. Error bars represent the standard error of the mean. A 1-way ANOVA with Tukey’s multiple comparisons test was used to compare the quantified protein amounts. For quantification of proteins, each blot was loaded with 5 standard proteins and three spore samples.

## Results

### Identifying *C. difficile* mutants with altered co-germinant requirements

In order to identify the receptor with which amino acid co-germinants interact, we used a strategy that was previously used to identify the bile acid germinant receptor (CspC) [37]. Although other strategies, such as Tn-seq, could be used to generate random mutations, most of these will result in germination null phenotypes and do not permit the screening of subtler phenotypes. As shown in Figure 1A, wild type *C. difficile* UK1 vegetative cells were exposed to EMS and recovered. Purified spores derived from the mutagenized bacteria were then germinated in buffer supplemented with 10 mM TA and 10 mM betaine. The structural difference between glycine and betaine is the presence of three methyl groups attached to the N-terminus rather than two hydrogen atoms. Because betaine is a glycine analog and does not stimulate spore germination when added with TA [48], we hypothesized that we could isolate change-of-function mutants that recognize betaine as a germinant or those that no longer require glycine as a co-germinant. Spores incubated in the presence of buffered TA and betaine were then plated and allowed to form spores. Potential mutants were enriched with this strategy 4 - 5 times before isolating colonies and screening for phenotypes.

Across several mutagenesis experiments, the most commonly-observed phenotypes were strains that did not require the co-germinant glycine to germinate and germinated in response to taurocholate only (TA-only). As shown in Figure 1B, wild type *C. difficile* UK1 spores required both TA and glycine to stimulate the release of CaDPA from the core. However, spores purified from isolates derived from separate EMS mutageneses released CaDPA in the presence of TA only (Figure 1C, 1D, and 1E). Importantly, though, these mutants still responded to glycine, the germination efficiency / rate increased upon glycine addition. To identify the mutation(s) that caused this phenotype, five different mutant strains (isolated from 4 independent EMS mutageneses) and a wild-type control were sent for genome re-sequencing. Surprisingly, when the sequences of the mutant strains were compared to the sequence of the wild-type parent, we identified SNPs common to all 5 mutants in the coding region or the promoter region of *yabG,* coding for a sporulation-specific protease (Table 1).

### Characterizing the function of *yabG*

To confirm that the TA-only phenotype is caused by a mutation in *yabG,* we inserted a group II intron into the *yabG* gene of *C. difficile* R20291 using TargeTron technology. Germination of the *C. difficile yabG::ermB* mutant (RS08) spores was compared to wild type using both OD_600_ and CaDPA release assays (Figure 2). As shown in Figure 2A, wild-type *C. difficile* R20291 spores required both TA and glycine in order to germinate. Interestingly, the RS08 mutant spores germinated in response to TA-only and this germination phenotype was not enhanced by the addition of glycine (in contrast to the phenotype of the EMS-mutant spores) (Figure 2B). We also tested L-alanine as a co-germinant instead of glycine (Figure S2), however, the spores did not show any enhancement in germination with the addition of L-alanine and germinated only in response to TA alone. When the mutation was complemented *in trans* by expression of *yabG* from a plasmid (pRS97), the spores again recognized glycine as a germinant (Figure 2C). The TA-only phenotype in the mutant spores was also confirmed by CaDPA release and compared to spores from both wild type and the complemented strain (Figure 2D, 2E, 2F).

**Figure 2.**
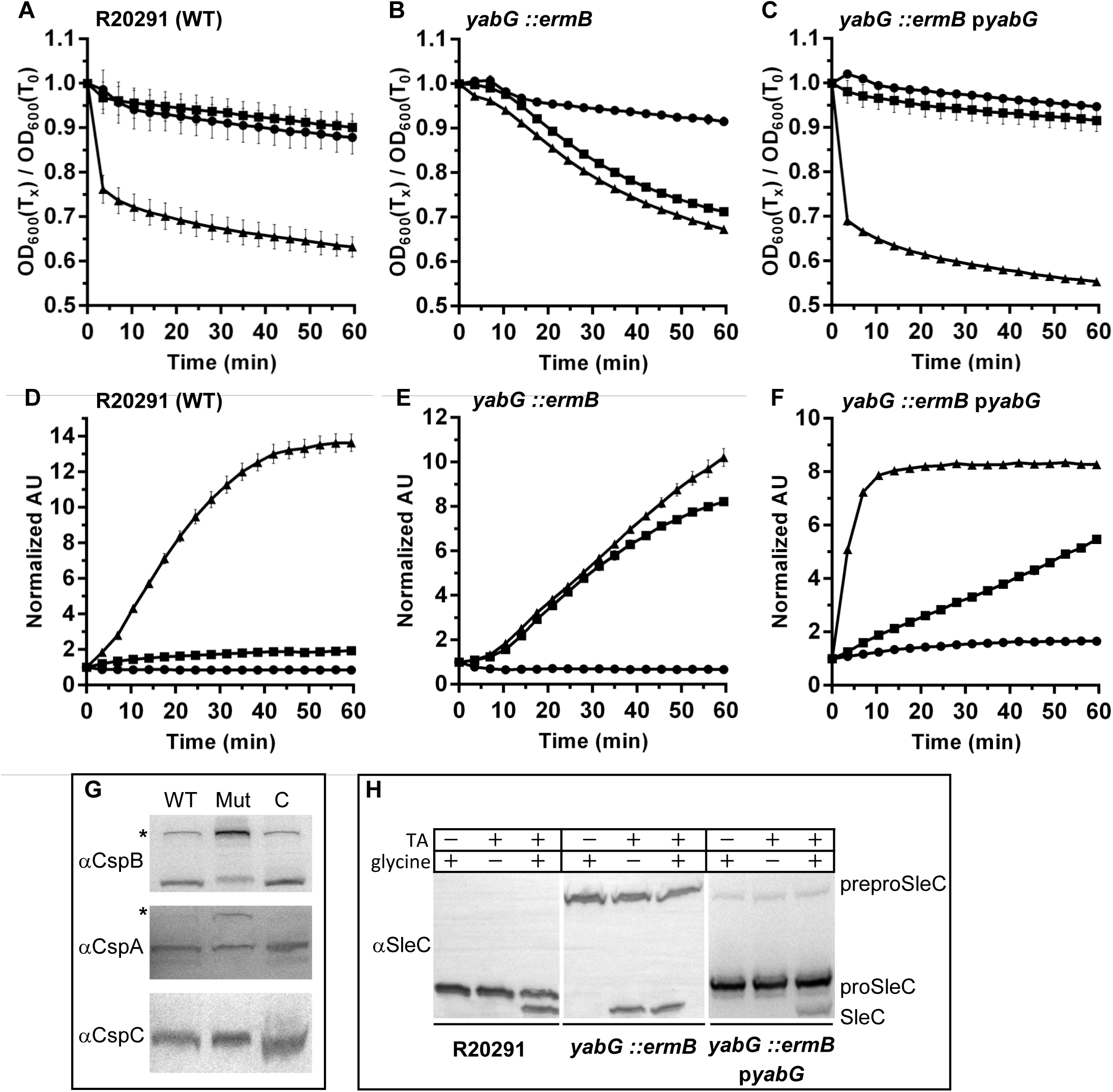
A mutation in *C. difficile yabG* results in spores that do not respond to amino acids as co-germinants. Spores derived from *C. difficile* R20291, *yabG::ermB* and *yabG::ermB* pRS97 (p*yabG*) strains were suspended in buffer supplemented with (●) 30 mM glycine or (■) 10 mM TA or (▲) 10 mM TA and 30 mM glycine. Germination was monitored at OD_600_ (A, B, C) or by the release of CaDPA in presence of 250 μM Tb^3+^ (D, E, F), respectively. Data points represent the averages from three, independent experiments and error bars represent the standard error of the mean. (G) Equal numbers of spores were extracted with NuPAGE buffer and separated by SDS-PAGE followed by immunoblotting with antisera specific for each listed protein. (H) Equal numbers of spores were incubated in 30 mM glycine or 10 mM TA or 10 mM TA and 30 mM glycine. Spores were then extracted and separated by SDS-PAGE. SleC activation was analyzed using antisera specific for the SleC protein (the SleC antibody detects the preproSleC, proSleC and activated SleC forms).

In prior work, a *yabG* mutant strain accumulated unprocessed CspBA into spores [42]. To confirm that the generated *yabG* mutant results in the accumulation of CspBA, we extracted spores derived from R20291, RS08 and the complemented strain and separated the extracted protein by SDS-PAGE. The separated protein was then detected using immunoblotting with CspB-specific antisera (Figure 2G). Although the blot showed that some CspBA is still remaining in both the wild type and complemented strains, the *yabG* mutant had mostly the unprocessed, CspBA form. Similarly, the CspA western blot showed that the *yabG* mutant had both the CspBA and CspA forms. The mutation in *yabG* did not appear to affect CspC incorporation into the spores although these results are not quantitative.

YabG also was to be required for the processing of preproSleC into proSleC [42]. Indeed, whereas the wild-type and complemented strains incorporated into spores the processed, proSleC, form, only preproSleC was incorporated into the RS08 strain (Figure 2H). When tested for the processing of SleC during germination, the wild-type and the complemented strains required both TA and glycine to activate SleC. However, *C. difficile* RS08 activated SleC in response to TA alone. These results confirm the TA-only phenotype observed in Figures 2B and 2E and suggest that a protein that is not processed in the *yabG* mutant strain is involved in germinant recognition or regulating the germinant specificity.

### Quantifying levels of CspB, CspC and SleC in spores from various strains

Because YabG is a sporulation specific protease, it is possible that the deletion of this protease might alter the amount of germination related proteins (*e.g.,* CspB, CspC, CspA or SleC) that are incorporated in the spore thereby providing the observed phenotype (i.e., increasing the abundance of the germinant receptors could lead to an increase in germinant sensitivity and loss of regulation). Using the previously described method to quantify the protein levels in *C. difficile* spores [33], we quantified the abundance of CspB (and CspBA), CspC and SleC in *C. difficile* RS08 (*yabG::ermB*) and compared them with the abundances in the wild-type and complemented strains (Figure 3).

**Figure 3.**
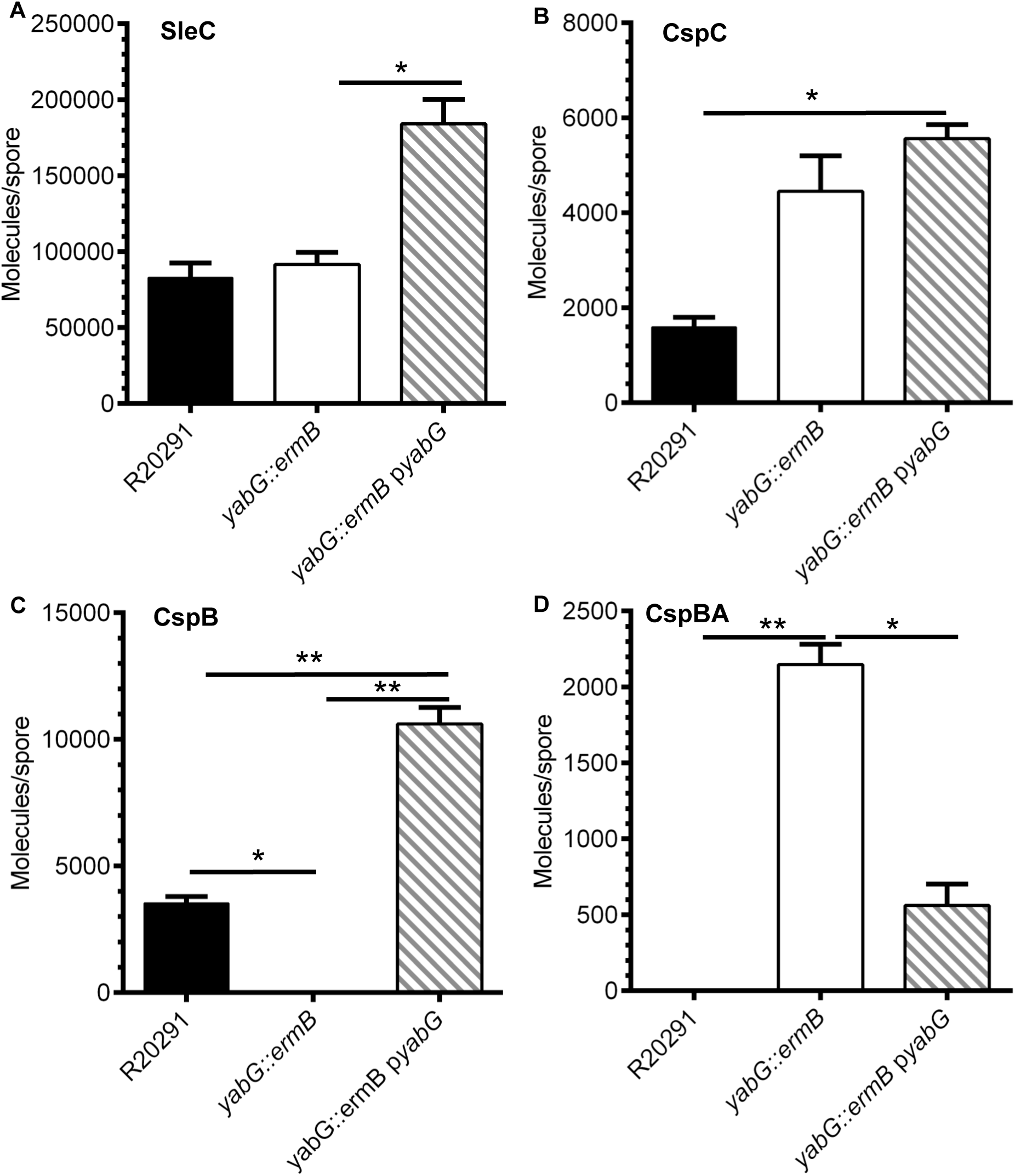
Quantifying the abundance of CspB, CspC and SleC in *C. difficile* spores. 2 × 10^9^ spores purified from *C. difficile* R20291, the *yabG* mutant and complemented strains and were extracted with NuPAGE buffer. Samples were separated by SDS-PAGE, transferred to low fluorescence PVDF membranes and blotted with antisera specific to the indicated proteins. The primary antibody was detected with a fluorescently conjugated secondary antibody and quantified as described in materials and methods. Quantified proteins are expressed as molecules / spore and are represented in a bar graph form. (A) SleC, (B) CspC, (C) CspB and (D) CspBA. The data presented represent the averages from three independent experiments and error bars represent the standard error of the mean. Statistical significance was determined using a 1 way ANOVA with Tukey’s multiple comparisons test (* p < 0.05; ** p < 0.01). *yabG::ermB* only incorporates the preproSleC form.

Introducing the *yabG* complementing plasmid resulted in significantly increased incorporation of proSleC and CspC into spores compared to the RS08 strain (Figure 3A; only the preproSleC form is incorporated into the RS08 strain) or the wild-type strain (Figure 3B), respectively. There were no statistical differences in the abundance of SleC or CspC between the wild-type and mutant strains. Importantly, there was a statistical difference in the abundance of CspB in all pair-wise comparisons of the wild type and complemented strain (there was no quantifiable CspB protein in spores derived from the RS08 strain; Figure 3C). Finally, spores derived from the RS08 strain had significantly more CspBA incorporated than did the wild-type or the complemented strains (Figure 3D). Because spores derived from the complemented strain had increased abundances of proSleC (Figure 3A), CspC (Figure 3B), and CspB (Figure 3C), but did not produce a TA-only phenotype, this suggests that increased abundance of these proteins is not the reason for the observed TA-only phenotype in the *yabG* mutant strain. Importantly, though, the *yabG* mutant accumulated much more CspBA into spores than did the wild-type or the complemented mutant strains (Figure 3D) and only accumulated the preproSleC form (Figure 2H). Therefore, we hypothesized that the presence of full-length CspBA and / or preproSleC could contribute to the observed TA-only phenotype.

### Deletions in the *cspBA* coding sequence lead to the observed TA-only phenotype.

Because CspB and CspC are already known to be involved in regulating *C. difficile* spore germination [37, 42–44, 46], we chose to first focus on the potential processing of CspBA by YabG. In prior work by Kevorkian *et. al.* [42], the CspBA processing site was hypothesized to occur at or near amino acid 548. To test this hypothesis, we deleted from the *C. difficile* CRG2359 (*C. difficile* R20291 Δ*pyrE*) chromosome 12 aa between CspB and CspA (*cspBA*_Δ548–560_) using *pyrE*-mediated allelic exchange. After confirmation of the engineered mutation in the CRG2359 genome, the *pyrE* gene was restored, and germination of the resulting strain was compared to *pyrE*-restored CRG2359 strain (*C. difficile* RS19; Figure S3A). We found that the CspBA_Δ548–560_ allele did not have any effect on germination (Figure S3B).

Next, to determine if deletions in the coding region in or between *cspB* and *cspA* affect spore germination, we deleted various regions within the *cspBA* gene in the CRG2359 strain (Figure 4A). The results of the germination phenotype in various deletions are shown in Figure 4. Deletion of 26 codons from the C-terminus of *cspB* (RS20; CspBA_Δ522–548_) did not affect spore germination (Figure 4C) when compared to the wild-type CRG2359 strain (Figure 4B). These results suggest that the C-terminus of CspB is not involved in generating the TA-only phenotype. Interestingly, deletion of 26 codons at the N-terminus of *cspA* (RS21; CspBA_Δ560–586_) resulted in spores that germinated in response to TA-only, after 60 minutes of incubation in the germination solution (Figure 4D). Though deletion of 25 codons in between the *cspB* and *cspA* coding sequences did not result in a TA-only phenotype (RS23; CspBA_Δ542–567_) (Figure 4E), extending the deleted region by another 9 codons into the surrounding region (RS24; CspBA_Δ537–571_) resulted in spores that germinated in response to TA-only, again after 60 minutes of incubation (Figure 4E). Importantly, the spores derived from the RS21 and RS24 strains still respond to glycine as co-germinant, despite also germinating in response to TA-only. Because the TA-only phenotype appeared to be enhanced as more *cspA* was deleted (RS21 to the RS24 strain), we predicted that a larger deletion might result in a phenotype similar to that observed in the *yabG* mutant (which does not recognize glycine as a co-germinant). We found that when 50 codons were deleted from the N-terminus of CspA (RS27) the spores no longer germinated (Figure 4F). These results were confirmed by analyzing the release of CaDPA from the germinating spores (Figure S4).

**Figure 4.**
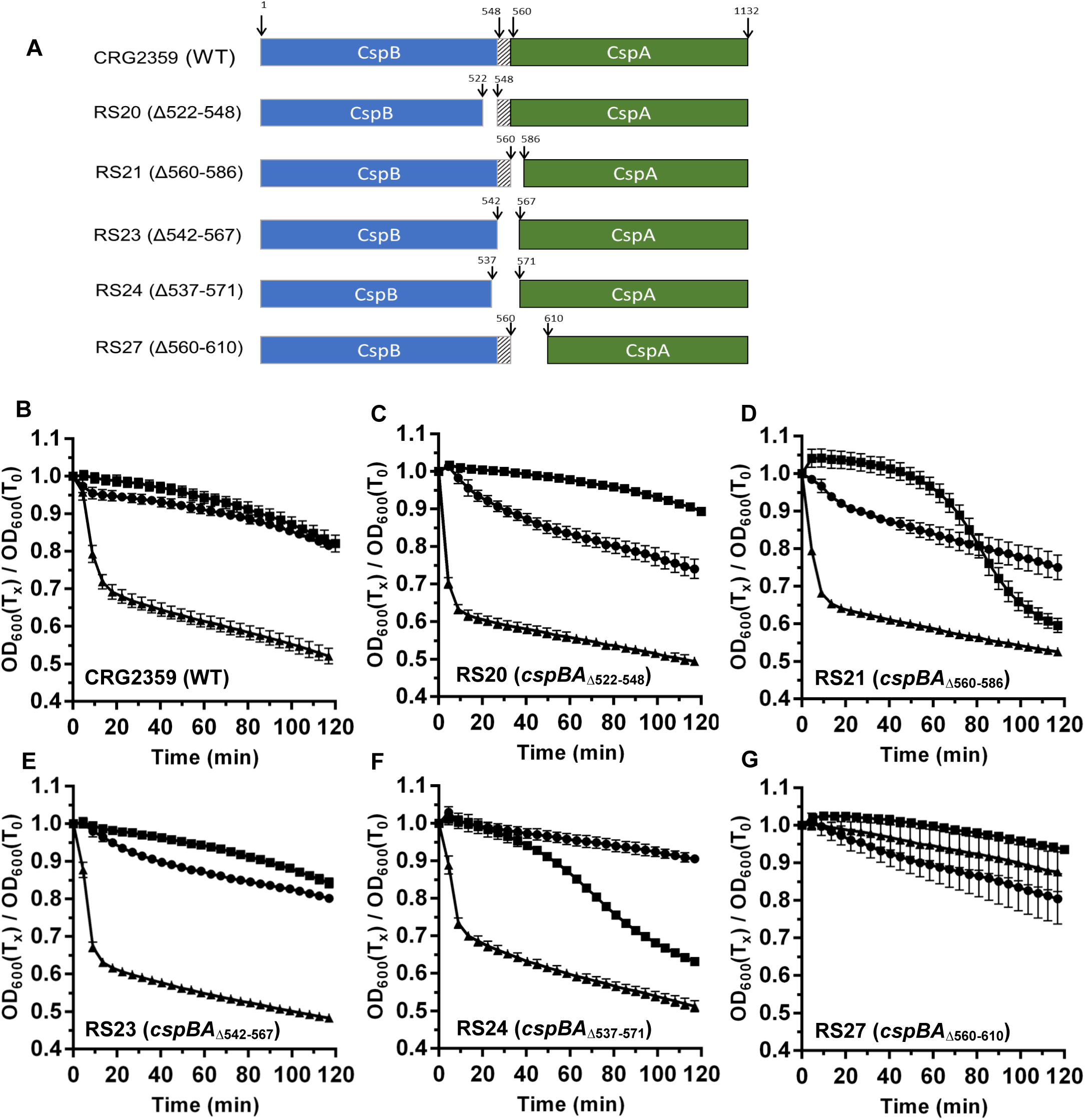
The N-terminus of *C. difficile* CspA is important for regulating germination in response to glycine. (A) Graphical representation of the various deletions introduced into *cspBA* compared to the parental, CRG2359; R20291Δ*pyrE* strain. Spore germination of the indicated strain was monitored at OD_600_ in buffer supplemented with (●) 30 mM glycine or (■) 10 mM TA or (▲) 10 mM TA and 30 mM glycine. (B) CRG2359, (C) RS20 (cspBA_Δ522–548_), (D) RS21 (*cspBA*_Δ560–586_), (E) RS23 (*cspBA*_Δ542–567_), (F) RS24 (*cspBA*_Δ537–571_) and (G) RS27 (*cspBA,*_Δ560–610_).

During construction of the deletion strains, we encountered significant difficulties in restoring the *pyrE* allele to wild type. To circumvent this obstacle, and to understand if *C. difficile* spore germination is affected by the *pyrE* deletion, we restored *pyrE* in the RS21 strain (CspBA_Δ560–586_, *pyrE*^+^; RS26) and compared with the RS19 strain (CRG2359 with restored *pyrE*). As shown in Figure S5A, *C. difficile* RS19 required both TA and glycine to germinate but the RS26 strain germinated in response to TA-only (Figure S5B). These results were confirmed by analyzing the release of CaDPA from the spore (Figure S5C and S5D). Finally, we analyzed the activation of proSleC to SleC in response to TA alone. Only the RS26 strain cleaved proSleC in response to TA alone (Figure S5E). These observations are identical to the observations made for the RS21 strain (Figure 4 and Figure S4) and indicate that the *pyrE* allele does not influence *C. difficile* spore germination in the context of these studies.

To confirm our observations that the strains germinated in response to TA alone, we analyzed by western blot the activation of SleC (Figure 5A). SleC activation in response to 10 mM TA only occurred in RS21 and RS24 while RS27 did not germinate in response to TA and glycine. The CspB western blot showed that CspBA was processed to CspB and CspA in all of the mutants, compared to wild type (Figure 5B). The CspC western blot did not reveal any differences between the wild type and mutant strains, except for the RS27 strain where no CspC was detected.

**Figure 5.**
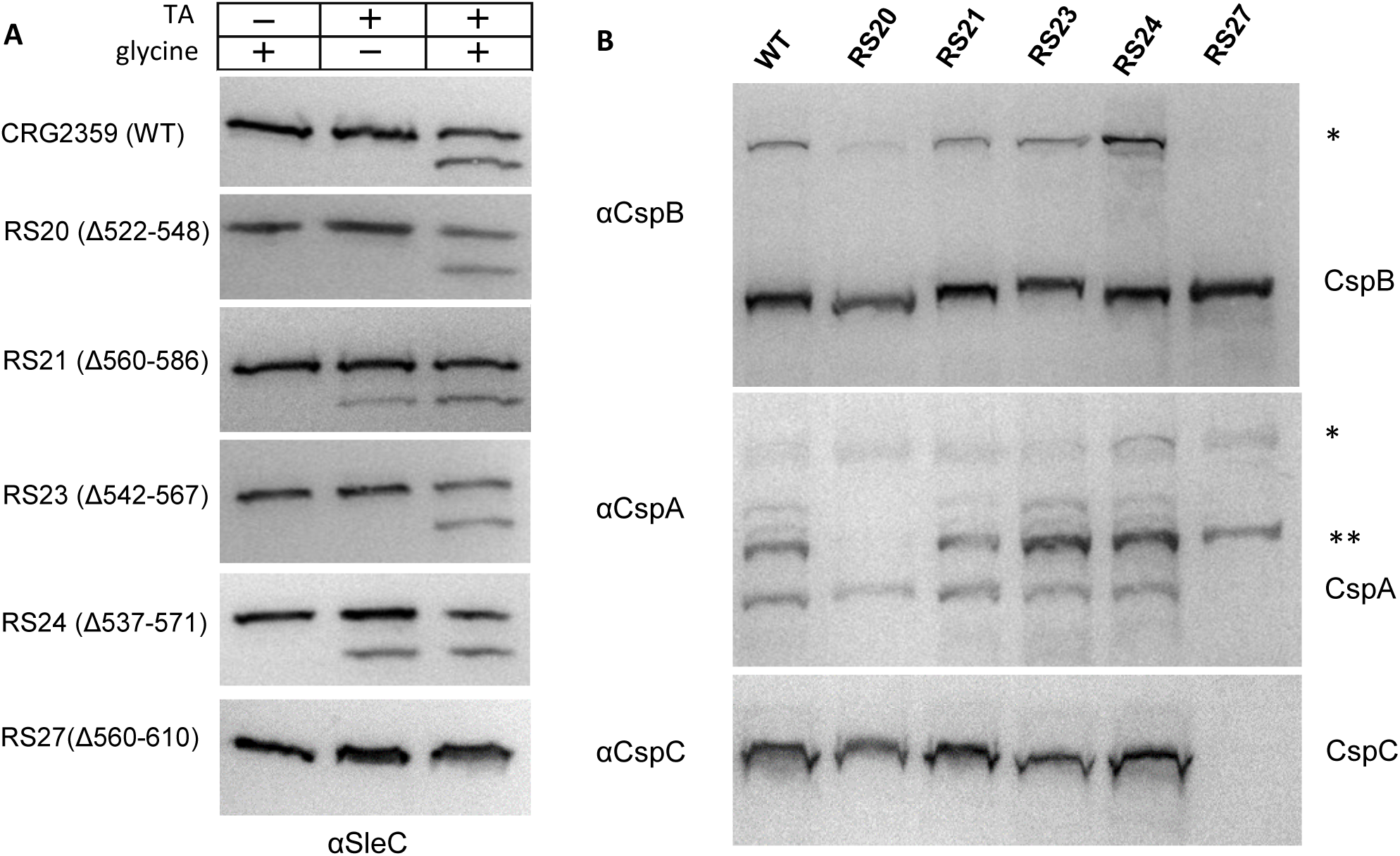
Comparing the effects of mutations in *C. difficile cspBA* on the incorporation and processing of CspB, CspA, CspC and proSleC. Spores derived from the indicated strains were extracted and separated as described in Figure 2. (A) SleC activation was measured in buffer supplemented with 30 mM glycine or 10 mM TA or both 10 mM TA and 30 mM glycine for 2 hours at 37 °C and (B) CspB, CspA, and CspC were detected as described in Figure 2 (*full length CspBA, ** alternative processing of CspA).

### Deletion of a hypothesized YabG cleavage site in CspA and preproSleC results in the TA-only phenotype

The data presented above suggests that the N-terminus of CspA is important for regulating spore germination. We had hypothesized that this region is processed by YabG but all the generated *cspA* alleles generated a protein that was processed into the CspB and CspA forms. One way to identify the YabG processing site in CspA is by pulldown of CspA from the spore extract and sequencing the CspA protein using mass spectrometry. Unfortunately, the CspA antibody was unable to immunoprecipitate CspA from the spores due to the quality of the antibody. However, we predicted that the YabG processing site in CspBA might be conserved in preproSleC. Instead of immunoprecipitating CspA, we immunoprecipitated proSleC from spore extracts derived from wild-type spores and *sleC* mutant spores (as a negative control) (Figure 6A). Using the sample from the proSleC pull down, we identied fragments by mass spectrometry of trypsin-digested proSleC (Figure 6B). In this experiment, the most N-terminal fragment identified began with glutamine followed by serine (an SR**<u>Q</u>**S sequence). When we compared this sequence to the protein sequence in CspA, we found that within the N-terminus of CspA there was a SRQS amino acid sequence that was encompassed within the deletions found in the RS21 strain (a strain that generated a TA-only phenotype).

**Figure 6.**
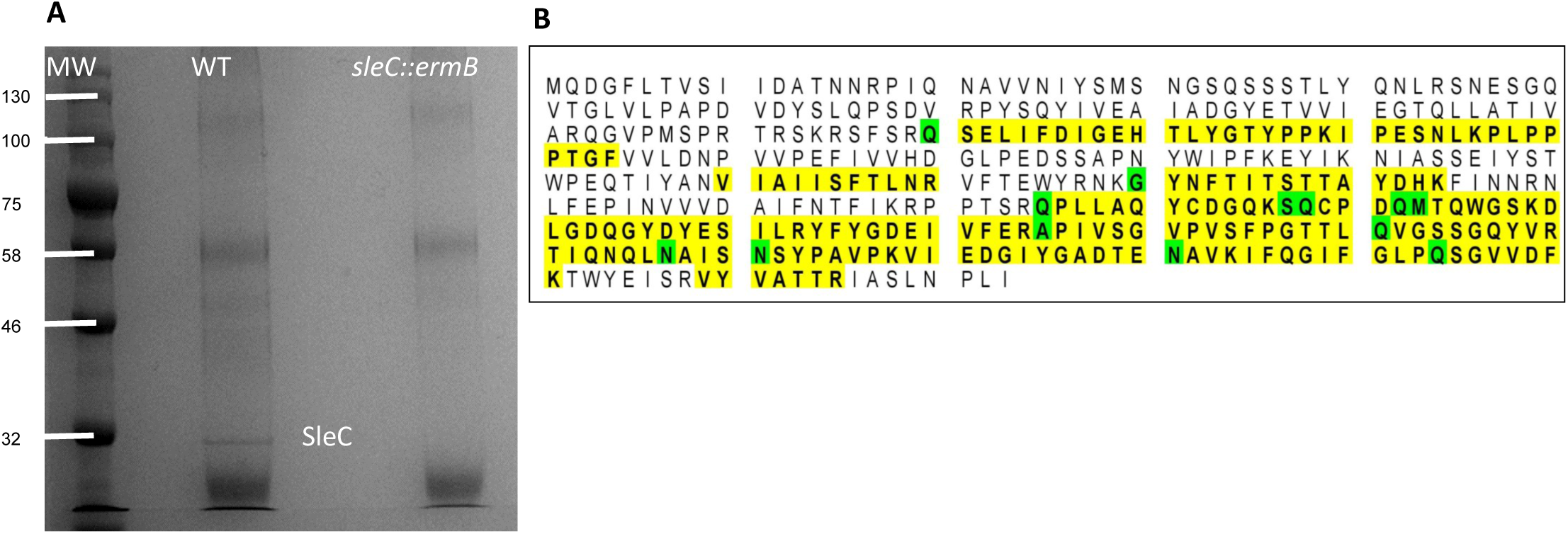
Immunoprecipitation of proSleC from *C. difficile* spores reveals a potential YabG cleavage sequence. (A) Spores derived from *C. difficile* R20291 (WT) and *C. difficile* RS10 (*sleC::ermB*) were disrupted by bead-beating and then proSleC was immunoprecipitated using SleC-specific antisera. Samples were separated by SDS-PAGE gel and stained with Commassie blue. The band corresponding to proSleC was excised from the gel and analyzed by peptide mass finger printing. (B) The sequence of preproSleC is listed. Yellow highlighted regions correspond to fragments that were detected by mass spectrometry and amino acids highlighted in gree indicate amino acids that are modified in the MS analysis (*i.e.,* oxidated, deaminated, acetylated or ammonia loss).

We deleted the nucleic acid sequence that encodes this SRQS motif in *sleC, cspBA,* or in both genes (Figure 7A). Surprisingly, the deletion of the SRQS site in CspA resulted in spores that germinated in response to TA-only within 40 minutes after germinant addition (Figure 7B). However, deletion of the SRQS motif within preproSleC did not affect spore germination (Figure 7C). When these two deletions were combined in the same strain, the spores had a TA-only phenotype similar to that of spores with a deletion in CspA alone (Figure 7D and 7A). Next, we confirmed these phenotypes by analyzing the release of CaDPA from the spore (Figure S6). Again, only when the *cspA*_ΔSRQS_ allele was incorporated into spores did the resulting strains release CaDPA in response to TA-only (Figure S6A and S6C); *sleC*_ΔSRQS_ spores required both TA and glycine to release CaDPA (Figure S6B).

**Figure 7.**
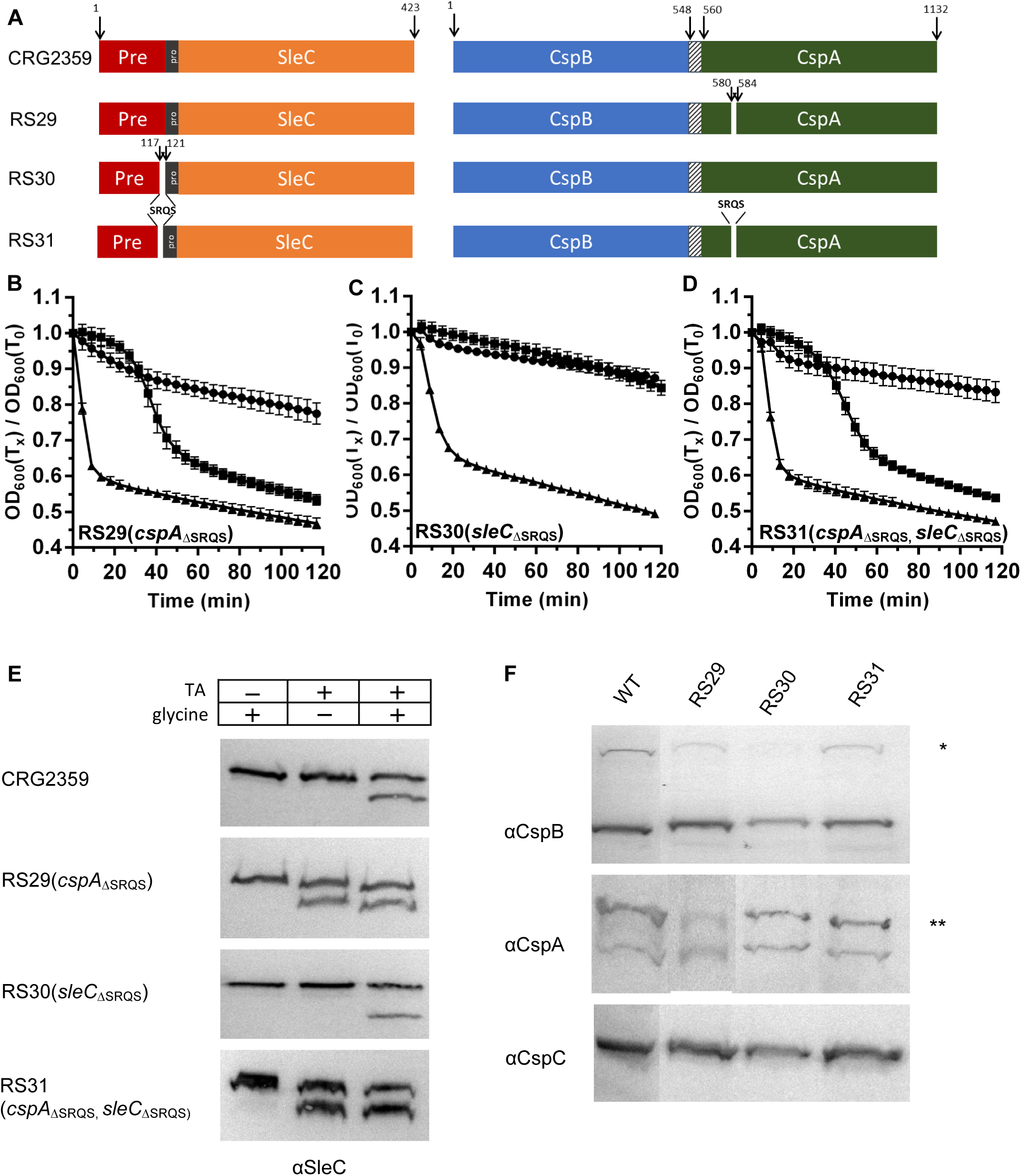
Deletion of the hypothesized YabG cleavage site in CspA results in a TA-only phenotype. (A) Graphical representation of the various deletions introduced into *cspBA* compared to the parental, CRG2359; R20291Δ*pyrE* strain. Spore germination of the indicated strain was monitored at OD_600_ in buffer supplemented with (●) 30 mM glycine or (■) 10 mM TA or (▲) 10 mM TA and 30 mM glycine. (B) RS29 (*cspA*_ΔSRQS_), (C) RS30 (*sleC*_ΔSRQS_), (D) RS31 (*cspA*_ΔSRQS_, *sleC*_ΔSRQS_). (E) Spores derived from the indicated strains were extracted and separated as described in Figure 2 and SleC activation was measured in buffer supplemented with 30 mM glycine or 10 mM TA or both 10 mM TA and 30 mM glycine for 2 hours at 37 °C. (F) CspB, CspA, and CspC were detected as described in Figure 2 (*full length CspBA, ** alternative processing of CspA).

We next analyzed the activation of SleC for these deletions. proSleC was activated in response to TA-only phenotype in the RS29 (*cspA*_ΔSRQS_) and the RS31 (*cspA*_ΔSRQS_ + *sleC*_ΔSRQS_) strains but not in the RS30 (*sleC*_ΔSRQS_) strain (Figure 7E). Interestingly, we also noticed that deletion of the SRQS sequence in the RS30 and RS31 strains did not result in the incorporation of full length preproSleC into the spores. Rather, preproSleC was still processed to a size consistent with a proSleC form, potentially suggesting that the SRQS sequence is not the YabG cleavage site or the presence of an alternative YabG processing site in preproSleC. There was no difference in the CspB, CspC and CspA incorporation in these SRQS deletion mutants compared to wild type (CRS2359; Figure 7F). Regardless, our results indicate that due to mis-processing of CspBA, or alterations within the *cspA* sequence, spores lose the requirement for an amino acid co-germinant during spore germination.

## Discussion

Previous studies on *C. difficile* spore germination have hypothesized the presence of an amino acid co-germinant receptor. We used EMS mutagenesis to screen for *C. difficile* mutants whose spores recognize betaine as a germinant. Excitingly, we isolated mutants that did not require an amino acid as a co-germinant, and these EMS-mutants germinated in response to TA-only. However, these mutants still recognized glycine as a co-germinant; the rate of germination increased upon the addition of glycine to the germination solution. Sequencing of these strains revealed common mutations in the *yabG* coding region or promoter region (Table 1). Thus, to confirm the TA-only phenotype observed in the EMS-generated mutants is due to the *yabG* allele, we created a TargeTron mutation in *yabG* of *C. difficile* R20291. In contrast to the EMS-generated mutants, the *yabG* mutant spores did not recognize glycine as a co-germinant (Figure 1C, 1D, 1E), and germinated in response to TA-only (Figure 2B, 2C). We hypothesize that because a small fraction of the CspBA pool in the EMS mutant strains is processed to CspB and CspA (Figure 2G), that the *yabG* alleles in these mutant strains generate less protein (for the promoter mutants) or less active protein (for mutants in the coding sequence). Under these conditions, the strains can still response to amino acids as co-germinants. However, in the TargeTron mutant, CspBA is unprocessed and the strain is unable to recognize amino acids. These phenotypes could be complemented by expressing *yabG in trans* from a plasmid. Oddly, we noticed that spores derived from the complemented strain released CaDPA in response to TA alone but this was not observed when germination was measured at OD_600_ (Figure 2F vs. 2C). We further analyzed this observation and found that the supposed CaDPA release observed in the complemented strain in the presence of TA alone was due the presence of Tb^3+^ in the assay (Tb^3+^ is absent from the germination solution in the OD_600_ assay). We are preparing a separate manuscript to report this in detail.

To understand if spores derived from the *yabG* mutant recognized other amino acids as co-germinants, we tested L-alanine. L-alanine is the second-best co-germinant during *C. difficile* spore germination with an EC_50_ value of 5 mM [32, 48]. Because *C. difficile* RS08 (*yabG::ermB*) spores did not respond to glycine or L-alanine (Figure S2), these results suggest that the protein(s) YabG processes is / are responsible for recognition of the various amino acids used as co-germinant(s). It is unclear if all of the amino acid co-germinants are recognized by a single protein or if each co-germinant / groups of co-germinants are recognized by unique proteins. However, our results suggest that glycine and other amino acids that trigger germination are recognized by a protein, or a set of proteins, that is processed by YabG.

YabG has been mostly studied in *B. subtilis.* In *B. subtilis* YabG is a sporulation-specific protease, and a *yabG* mutation causes alterations in the coat proteins of *B. subtilis* spores. The orthologues of most *B. subtilis* YabG target proteins are absent in *C. difficile* (*e.g.,* CotT, YeeK, YxeE, CotF, YrbA) [49–51]. In a previous report, the process of CspBA and preproSleC during *C. difficile* sporulation was shown to be YabG-dependent [42]. However, germination was not significantly altered in the mutant spores compared to wild type spores (germination efficiency decreased from 1 to 0.8 in mutants, when analyzing CFU counts) [42]. Importantly, though, germination efficiency was only tested on BHIS-TA agar medium and not in the presence of TA alone [42].

In prior work by Kevorkian *et. al.* [42], the authors suggested that the CspBA processing site is near-amino acid 548 and is encoded by a linker DNA sequence between *cspB* and *cspA.* To test this, we used allelic exchange to delete the codons encoding amino acids 542–567 regions in *cspBA* (RS23). When the germination phenotype of the RS23 strain was compared with CRG2359, we did not observe a TA-only phenotype (Figure 4A and 4E) and, importantly, CspBA was still efficiently processed (Figure 5). These results suggest that the predicted 548–560 region as the YabG-dependent processing site is not accurate, or that when the YabG processing site is removed, YabG cleaves the CspBA protein at an alternate site.

Unlike spores derived from the *yabG* mutant, which did not recognize amino acids as co-germinants, the deletions that were engineered in the N-terminus in CspA [*e.g.,* CspBA_Δ560–586_ (RS21) or CspBA_ΔSRQS_ (RS29)] still responded to glycine as a co-germinant. In order to recapitulate the *yabG* mutant phenotype, we hypothesized that deletion of a larger portion in the N-terminus of CspA might result in a CspA allele that loses the recognition of amino acids as co-germinants. When 50 codons were deleted from the region of CspA encoding the N-terminus (CspBA_Δ560–610_; RS27), the spores no longer germinated (Figure 4G) and no SleC was activated in any of the tested conditions (Figure 5). Moreover, we did not observe functional CspA in the spores derived from the RS27 (although there was a band corresponding to unprocessed CspA similar to what was observed in the other deletion and wild-type strains). Prior work on CspA demonstrated that deletion of CspA results in spores that do not incorporate CspC [42, 43]. Here, we found that the 50 amino acids at N-terminus of CspA are important for the incorporation of CspC into spores, probably because CspA is non-functional without the N-terminal domain or the mutant lacks a localization signal that directs the processed CspA into the spore.

Using immunoprecipitation and mass spectrometry of the immunoprecipitated protein, we identified four amino acids in SleC with identity in CspBA. When this sequence was deleted from CspA (*cspBA*_Δ580–584_), we observed a TA-only phenotype in the spores suggesting that the SRQS region might be important for CspA activity. Both RS21 (*cspBA*_Δ560–586_) and RS24 (*cspBA*_Δ537–571_) strains generated spores with TA-only phenotypes similar to the SRQS deletion in CspA (RS29; *cspBA*_Δ580–584_). However, only the RS21 strain has the deletion of SRQS region while the SRQS motif is still present in the RS24 strain. Importantly, the SRQS deletion in preproSleC resulted in spores with no TA-only phenotype and the western blot analysis (Figure 7) showed that deletion of the SRQS site in preproSleC resulted in incorporation of the processed, proSleC, into the spore instead of preproSleC form. This indicates that the SRQS sequence is not the YabG processing site in preproSleC (and, potentially CspBA), or that the deletion of the SRQS sequence results in YabG shifting its processing site in preproSleC to a region near the SRQS sequence. Further experiments are required to identify the YabG cleavage sites in spore proteins and how this processing is affected when its recognition site is deleted.

In a recent review on *C. difficile* spore germination [13], the authors build upon a hypothesized model for spore germination [10] and propose a new, “lock and key” model for *C. difficile* spore germination. In the first model, the germinosome complex composed of CspC, CspA, CspB, and proSleC are anchored to the outer spore membrane by GerS (a lipoprotein that is required for spore germination [52] - see below) [10, 13]. Upon binding of a germinant (TA with glycine or Ca^2+^), CspC and CspA are released from this germinosome complex and CspB becomes free to activate proSleC to degrade the cortex. Subsequently, CaDPA is released from the spore core by a mechanosensing mechanism [26, 45]. In the second model, CspA and CspC are localized in the coat layer, where TA can bind to CspC. Activation of CspC leads to the transport of glycine and Ca^2+^ through the outer membrane, by an unknown protein, to the cortex where CspB is held inactive in a complex with GerS and proSleC. In the absence of calcium, glycine is transported to the inner membrane where it activates the release of Ca^2+^ from the core through another unknown process. The released Ca^2+^ traffics to CspB to activate its protease activity. In the presence of calcium and glycine, both are transported in and Ca^2+^ activates CspB. When glycine and calcium bind to CspB, CspB can activate proSleC to degrade the cortex [13].

Our results support the first model whereby CspB, CspA, and CspC are in a germinosome complex – similar to the germinosome complex found in *B. subtilis* [53]. Recent studies on GerS have shown that GerS likely does not form a complex with other germination proteins [52, 54]. Rather, it is required to generate cortex-specific modifications in the spore and thus, important for germination of the spore because the SleC cortex lytic enzyme depends on cortex-specific modifications to degrade the cortex layer efficiently [54]. Based upon prior work from our lab, proSleC is unlikely to be part of this complex because SleC is three to four times more abundant than CspB or CspC, depending upon the strain analyzed [33].

The processing of CspBA depends on the YabG protease and *yabG*-mutant spores incorporate mostly the full length CspBA protein. These spores do not recognize amino acids as co-germinants and germinate in response to TA-only (Figure 2B). We hypothesize that CspC (functioning as the bile acid germinant receptor) and CspA (functioning as the amino acid co-germinant receptor) inhibit CspB activity within dormant spores (Figure 8). These two pseudoproteases would regulate the activity of CspB so that it does not prematurely activate proSleC, potentially similar to how other pseudoproteases / pseudokinases regulate activity of their cognate proteins [55–59]. Because *yabG* mutant spores package full length CspBA, where CspA is tethered to CspB, CspC alone prevents CspB from cleaving proSleC into its active form, and TA might dislodge CspC from CspB (this is consistent with our prior publication that indicates that CspC may have an inhibitory activity during spore germination [33]). Moreover, our data suggest that the N-terminus of CspA might be important for formation of this hypothesized complex with CspC and / or CspB. When portions of the CspA N-terminus are deleted, the binding of CspA to the complex might become unstable causing CspA to randomly disassociate from the complex. This would result in TA alone stimulating germination by disassociating CspC from CspB. Once CspB is free from the complex, it could then activate many proSleC proteins to maximize the germination process. Further work is needed to test the biochemical implications of this hypothesis.

**Figure 8.**
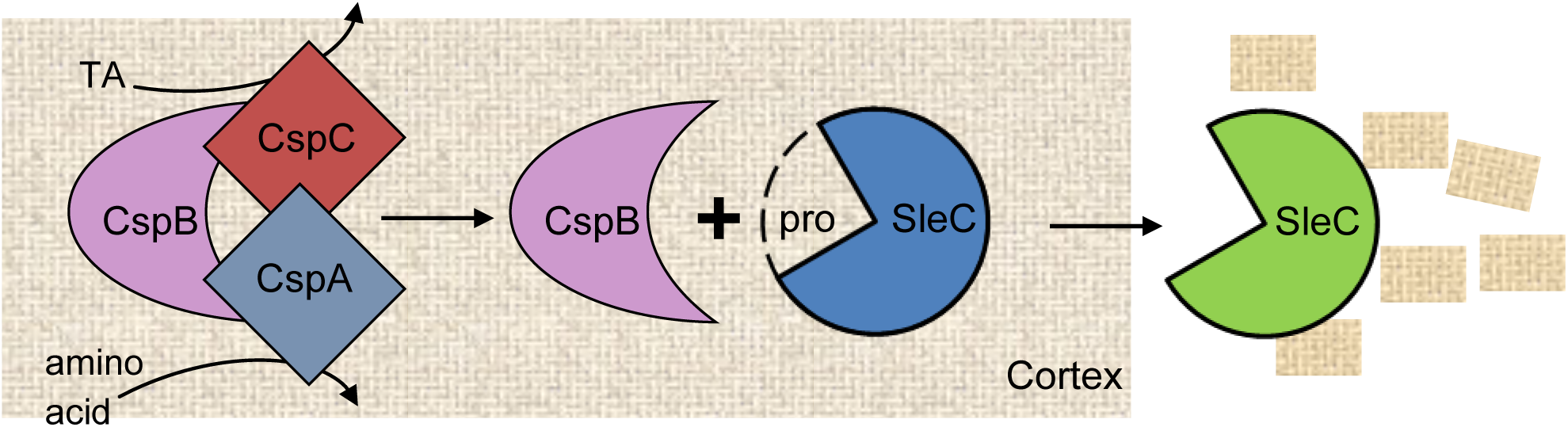
Model for *C. difficile* spore germination. In our working model for *C. difficile* spore germination, CspB protease activity is inhibited by two pseudoproteases, CspC (the bile acid germinant receptor; [37]) and CspA (the amino acid germinant receptor; this study). Interaction of CspC with cholic acid derivatives [31] and the interaction of CspA with amino acids [32, 60] results in these two proteins disassociating from CspB. Subsequently, CspB cleaves the inhibitory pro-domain from proSleC thereby activating SleC’s cortex degrading activity.

## Acknowledgments

We thank members of the Sorg lab for critical reading of this manuscript and their helpful comments during this study. We also thank members of Dr. Leif Smith’s laboratory at Texas A&M University for helpful comments during preparation of this manuscript.

